# Ten Simple Rules for Taking Advantage of git and GitHub

**DOI:** 10.1101/048744

**Authors:** Yasset Perez-Riverol, Laurent Gatto, Rui Wang, Timo Sachsenberg, Julian Uszkoreit, Felipe da Veiga Leprevost, Christian Fufezan, Tobias Ternent, Stephen J. Eglen, Daniel S. Katz, Tom J Pollard, Alexander Konovalov, Robert M. Flight, Kai Blin, Juan Antonio Vizcaino

## Abstract

A ‘Ten Simple Rules’ guide to git and GitHub. We describe and provide examples on how to use these software to track projects, as users, teams and organizations. We document collaborative development using branching and forking, interaction between collaborators using issues and continuous integration and automation using, for example, Travis CI and codevoc. We also describe dissemination and social aspects of GitHub such as GitHub pages, following and watching repositories, and give advice on how to make code citable.

## Introduction

Bioinformatics is a broad discipline in which one common denominator is the need to produce and/or use software that can be applied to biological data in different contexts. To enable and ensure the replicability and traceability of scientific claims, it is essential that the scientific publication, the corresponding datasets, and the data analysis are made publicly available [1, 2]. All software used for the analysis should be either carefully documented (e.g., for commercial software) or better, openly shared and directly accessible to others [3, 4]. The rise of openly available software and source code alongside concomitant collaborative development is facilitated by the existence of several code repository services such as SourceForge (http://sourceforge.net/), Bitbucket (https://bitbucket.org/), GitLab (https://about.gitlab.com/), and GitHub (https://github.com/), among others. These resources are also essential for collaborative software projects, since they enable the organization and sharing of programming tasks between different remote contributors. Here, we introduce the main features of GitHub, a popular web-based platform that offers a free and integrated environment for hosting the source code, documentation, and project-related web content for open source projects. GitHub also offers paid plans for private repositories (see Box 2) for individuals and businesses, as well as free plans including private repositories for research and educational use.

GitHub relies, at its core, on the well-known and open source version control system git, originally designed by Linus Torvalds for the development of the Linux kernel, and now developed and maintained by the git community (https://github.com/git). One reason for GitHub’s success is that it offers more than a simple source code hosting service [5, 6]. It provides developers and researchers with a dynamic and collaborative environment, often referred to as a social coding platform, that supports peer review, commenting and discussion [7]. A diverse range of efforts, ranging from individual to large bioinformatics projects, laboratory repositories, as well as global collaborations have found GitHub to be a productive place to share code, ideas and collaborate (see Table 1).

**Table 1.**
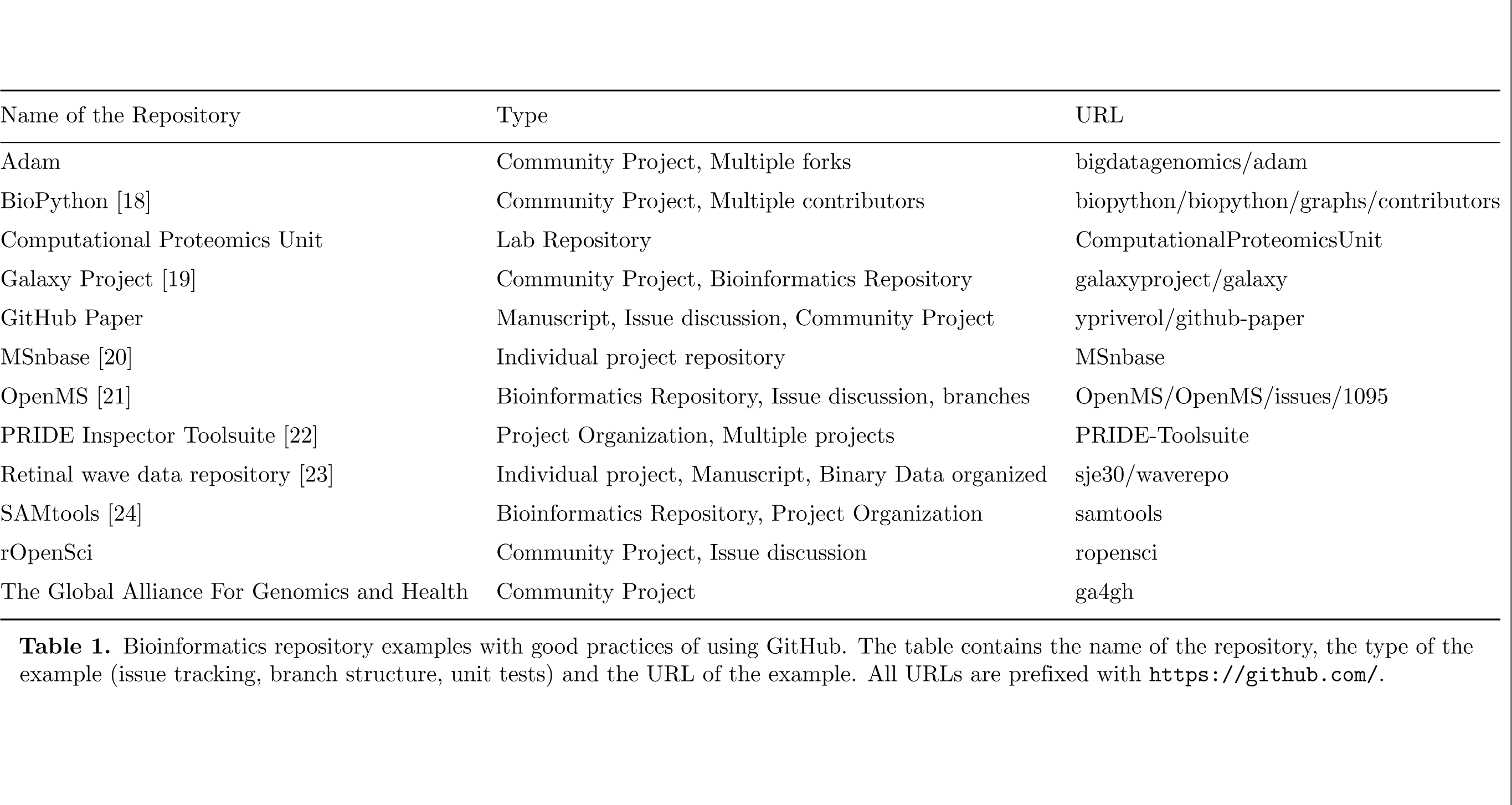
Bioinformatics repository examples with good practices of using GitHub. The table contains the name of the repository, the type of the example (issue tracking, branch structure, unit tests) and the URL of the example. All URLs are prefixed with https://github.com/.

Some of the recommendations outlined below are broadly applicable to repository hosting services. However our main aim is to highlight specific GitHub features. We provide a set of recommendations that we believe will help the reader to take full advantage of GitHub’s features for managing and promoting projects in bioinformatics as well as in many other research domains. The recommendations are ordered to reflect a typical development process: learning git and GitHub basics, collaboration, use of branches and pull requests, labeling and tagging of code snapshots, tracking project bugs and enhancements using issues, and dissemination of the final results.

## Rule 1. Use GitHub to track your projects

The backbone of GitHub is the distributed version control system *git*. Every change, from fixing a typo to a complete redesign of the software, is tracked and uniquely identified. While git has a complex set of commands and can be used for rather complex operations, learning to apply the basics requires only a handful of new concepts and commands, and will provide a solid ground to efficiently track code and related content for research projects. Many introductory and detailed tutorials are available (see Table 2 below for a few examples). In particular, we recommend *A Quick Introduction to Version Control with Git and GitHub* by Blischak *et al.* [5].

**Table 2.** Online courses, tutorials and workshops about GitHub and Git for scientists.

| Name of the material | URL |
| --- | --- |
| `git help` and `git help -a` | Document, installed with git |
| Karl Broman's git/github guide | http://kbroman.org/github_tutorial/ |
| Version Control with GitVersion Control with Git | http://swcarpentry.github.io/git-novice/ |
| Introduction to Git | http://git-scm.com/book/ch1-3.html |
| Github Training | https://training.github.com/ |
| Github Guides | https://guides.github.com/ |
| Good Resources for Learning Git and GitHub | https://help.github.com/articles/good-resources-for-learning-git-and-github/ |
| Software Carpentry: Version Control with Git | http://swcarpentry.github.io/git-novice/ |

In a nutshell, initialising a (local) repository (often abbreviated *repo*) marks a directory as one to be tracked (Fig. 1). All or parts of its content can be added explicitly to the list of files to track.

**Figure 1.**
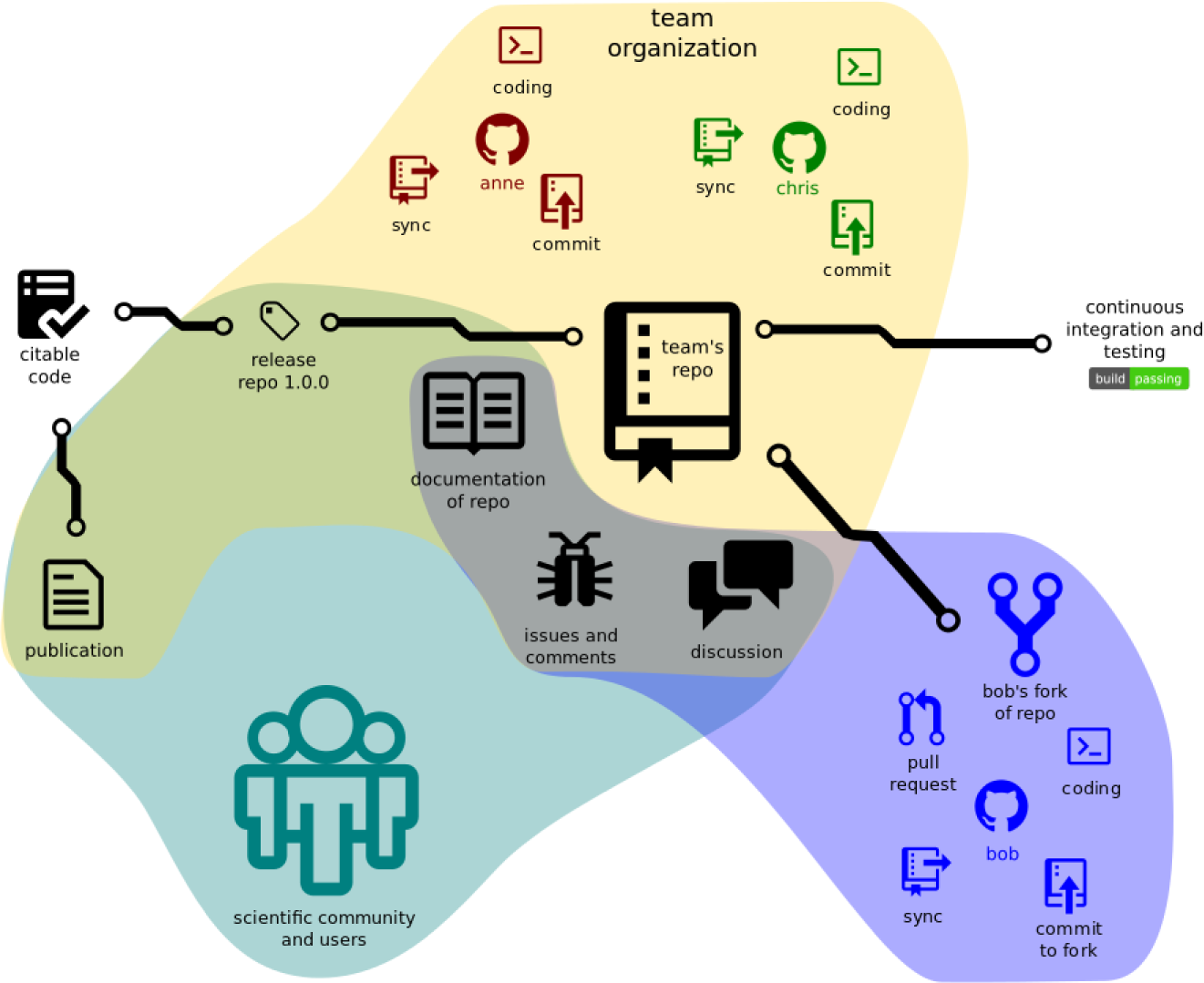
The structure of a GitHub-based project illustrating project structure and interactions with the community.

~~~
cd project ## move into directory to be tracked
~~~

~~~
git init ## initialise local repository
~~~

~~~
## add individual files such as project description, reports, source code
~~~

~~~
git add README project.md code.R
~~~

~~~
git commit -m “initial commit” ## saves the current local snapshot
~~~

Subsequently, every change to the tracked files, once committed, will be recorded as a new revision, or *snapshot*, uniquely identifying the changes in all the modified files. Git is remarkably effective and efficient in archiving the complete history of a project by, among other things, storing only the differences between files.

In addition to local copies of the repository, it is straightforward to create remote repositories on GitHub (called origin, with default branch master - see below) using the web interface, and then synchronize local and remote repositories.

~~~
git push origin master ## push local changes to the remote repository
~~~

~~~
git pull origin master ## pull remote changes into the local repository
~~~

Following Tony Rossini’s advice in 2005 to “commit early, commit often, and commit in a repository from which we can easily roll-back your mistakes”, one can organise their work in small incremental changes. At any time it is possible to go back to a previous version. In larger projects, multiple users are able to work on the same remote repository, with all contributions being recorded, restorable and attributed to the author.

Users usually track source code, text files, images, and small data files inside their repositories, and generally do not track derived files such as build logs or compiled binaries. And while the majority of GitHub repositories are used for software development, users can also keep text documents such as analysis reports and manuscripts (see, for example, the repository for this manuscript at https://github.com/ypriverol/github-paper).

Due to its distributed design, each up-to-date local git repository is an entire exact historical copy of everything that was committed - file changes, commit message logs, etc. These copies act as independent backups as well, present on each user’s storage device. Git can be considered to be fault-tolerant because of this, which is a win over centralized version control systems. If the remote GitHub server is unavailable, collaboration and work can continue between users, as opposed to centralized alternatives.

The web interface offered by GitHub provides friendly tools to perform many basic operations and a gentle introduction to a more rich and complex set of functionalities. Various graphical user-interface driven clients for managing git and GitHub repositories are also available (https://www.git-scm.com/downloads/guis). Many editors and development environments such as, for example, the popular RStudio editor (https://www.rstudio.com/) for the R programming language [8], directly integrate with code versioning using git and GitHub. In addition, for remote git repositories, GitHub provides its own features that will be described in subsequent rules (Fig. 1).

### Box 1

Using GitHub, or any similar versioning/tracking system is not a replacement for good project management; it is an extension, an improvement for good project and file managing (see for example [9]). One practical consideration when using GitHub, for example, is dealing with large binary files. Binary files such as images, videos, executable files, or many raw data used in bioinformatics, are stored as a single large entity in git. As a result, every change, even if minimal, leads to a complete new copy of the file in the repository, producing large size increments and the inability to search (see https://help.github.com/articles/searching-code/) and compare file content across revisions. Git offers a Large File Storage (LFS) module (https://git-lfs.github.com/) that replaces such large files with pointers, while the large binary file can be stored remotely, which results in small and faster repositories. Git LFS is also supported by GitHub, albeit with a space quota or for a fee, to retain your usual GitHub workflow (https://help.github.com/categories/managing-large-files/) (Supplementary File S1, Section 1).

### Box 2

By default, GitHub repositories are freely visible to all. Many projects decide to share their work publicly and openly from the start of the project, in order to attract visibility and to benefit from contributions from the community early on. Some other groups prefer to work privately on projects until they are ready to share their work. Private repositories ensure that work is hidden but also limit collaborations to just those users that are given access to the repository. These repositories can then be made public at a later stage, such as, for example, upon submission, acceptance, or publication of corresponding journal articles. In some cases, when the collaboration was exclusively meant to be private, some repositories might never be made publicly accessible.

### Box 3

Every repository should ideally have the following three files. The first, and arguably most important file in a repository is a LICENCE file (see also Rule 8), that clearly defines the permissions and restrictions attached to the code and other files in your repository. The second important file is a README file, which provides, for example, a short description of the project, a quick start guide, information on how to contribute, a TODO list, and links to additional documentation. Such README files are typically written in markdown, a simple markup language that is automatically rendered on GitHub. Finally, a CITATION file to the repository informs your users how to cite and credit your project.

## Rule 2. GitHub for single users, teams and organizations

Public projects on GitHub are visible to everyone, but write permission, i.e., the ability to directly modify the content of a repository, needs to be granted explicitly. As a repository owner, you can grant this right to other GitHub users. In addition to being owned by users, repositories can also be created and managed as part of teams and organizations.

Project managers can structure projects to manage permissions at different levels: users, teams and organizations. Users are the central element of GitHub, as in any other social network. Every user has a profile listing their GitHub projects and activities, which can optionally be populated with personal information including name, e-mail address, image, and webpage. To stay up to date with the activity of other users, one can *follow* their accounts (see also Rule 10). Collaboration can be achieved by simply adding a trusted *Collaborator*, thereby granting write access.

However, development in large projects is usually done by teams of people, within a larger organization. GitHub organizations are a great way to manage team-based access permissions for the individual projects of institutes, research labs, and large open source projects that need multiple owners and administrators (Fig. 1). We recommend that you, as an individual researcher, make your profile visible to other users and display all of the projects and organisations you are working in.

## Rule 3. Developing and collaborating on new features: branching and forking

Anyone with a GitHub account can *fork* any repository they have access to. This will create a complete copy of the content of the repository, while retaining a link to the original ‘upstream’ version. One can then start working on the same code base in one’s own fork (https://help.github.com/articles/fork-a-repo/) under their username (see, for example, https://github.com/ypriverol/github-paper/network/members for this work) or organization (see Rule 2). Forking a repository allows users to freely experiment with changes without affecting the original project and forms the basis of social coding. It allows anyone to develop and test novel features with existing code and offers the possibility of contributing novel features, bug fixes, and improvements to documentation back into the original upstream project (requested by opening an *pull request*) repository and becoming a contributor. Forking a repository and providing pull requests constitutes a simple method for collaboration inside loosely defined teams and over more formal organizational boundaries, with the original repository owner(s) retaining control over which external contributions are accepted. Once a pull request is opened for review and discussion, it usually results in additional insights and increased code quality [7].

Many contributors can work on the same repository at the same time without running into edit conflicts. There are multiple strategies for this, and the most common way is to use git *branches* to separate different lines of development. Active development is often performed on a development branch and stable versions, i.e., those used for a software release, are kept in a master or release branch (see for example https://github.com/OpenMS/OpenMS/branches). In practice, developers often work concurrently on one or several features or improvements. To keep commits of the different features logically separated, distinct branches are typically used. Later, when development is complete and verified to work (i.e., none of the tests fail, see Rule 5), new features can be merged back into the development line or master branch. In addition, one can always pull the currently up-to-date master branch into a feature branch, to adapt the feature to the changes in the master branch.

When developing different features in parallel, there is a risk of applying incompatible changes in different branches/forks; these are said to become *out of sync*. Branches are just short-term departures from master. If you pull frequently, you will keep your copy of the repository up to date, and you will have the opportunity to merge your changed code with others’ contributors, ideally without requiring you to manually address conflicts to bring the branches in sync again.

## Rule 4. Naming branches and commits: tags and semantic versions

Tags can be used to label versions during the development process. Version numbering should follow ‘semantic versioning’ practice, with the format X.Y.Z, with X being the major, Y the minor, and Z the patch version of the release, including possible meta information, as described in http://semver.org/. This semantic versioning scheme provides users with coherent version numbers that document the extent (bug fixes or new functionality) and backwards compatibility of new releases. Correct labeling allows developers and users to easily recover older versions, compare them, or simply use them to reproduce results described in publications (see Rule 8). This approach also help to define a coherent software publication strategy.

## Rule 5: Let GitHub do some tasks for you: integrate

The first rule of software development is that the code needs to be ready to use as soon as possible [10], to remain so during development, and that it should be well-documented and tested. In 2005, Martin Fowler defined the basic principles for continuous integration in software development [11]. These principles have become the main reference for best practices in continuous integration, providing the framework needed to deploy software, and in some way, also data. In addition to mere error-free execution, dedicated code testing is aimed at detecting possible bugs introduced by new features, or changes in the code or dependencies, as well as detecting wrong results, often known as *logic errors*, where the source code produces a different result than what was intended. Continuous integration provides a way to automatically and systematically run a series of tests to check integrity and performance of code, a task that can be automated through GitHub.

GitHub offers a set of *hooks* (automatically executed scripts) that are run after each push to a repository, making it easier to follow the basic principles of continuous integration. The GitHub web hooks allows third-party platforms to access and interact with a GitHub repository and thus to automate post-processing tasks. Continuous integration can be achieved by *Travis CI* (https://travis-ci.org), a hosted continued integration platform that is free for all open source projects. Travis CI builds and tests the source code using a plethora of options such as different platforms and interpreter versions (Supplementary File S1, Section 2). In addition, it offers notifications that allow your team and contributors to know if the new changes work, and to prevent the introduction of errors in the code (for instance, when merging pull requests), making the repository always ready to use.

## Rule 6: Let GitHub do more tasks for you: automate

More than just code compilation and testing can be integrated into your software project: GitHub hooks can be used to automate numerous tasks to help improve the overall quality of your project. An important complement to successful test completion is to demonstrate that the tests sufficiently cover the existing code base. For this, the integration of *Codecov* is recommended (https://codecov.io). This service will report how much of the code base and which lines of code are being executed as part of your code tests. The Bioconductor project, for example, highly recommends that packages implement unit testing (Supplementary File S1, Section 2) to support developers in their package development and maintenance (http://bioconductor.org/developers/unitTesting-guidelines/), and systematically tests the coverage of all of its packages (https://codecov.io/github/Bioconductor-mirror/). One might also consider generating the documentation upon code/documentation modification (Supplementary File S1, Section 3). This implies that your projects provide comprehensive documentation so others can understand and contribute back to them. For Python or C/C++ code, automatic documentation generation can be done using sphinx (http://sphinx-doc.org/) and subsequently integrated into GitHub using “Read the Docs” (https://readthedocs.org/). All of these platforms will create reports and badges (sometimes called shields) that can be included on your GitHub project page, helping to demonstrate that the content is of high quality and well-maintained.

## Rule 7. Use GitHub to openly and collaboratively discuss, address and close issues

GitHub *issues* are a great way to keep track of bugs, tasks, feature requests, and enhancements. While classical issue trackers are primarily intended to be used as bug trackers, in contrast, GitHub issue trackers follow a different philosophy: each tracker has its own section in every repository and can be used to trace bugs, new ideas, and enhancements, by using a powerful tagging system. *Issues* main focus is on promoting collaboration, providing context by using cross-references.

Raising an issue does not require lengthy forms to be completed. It only requires a title, and preferably at least a short description. Issues have very clear formatting, and provide space for optional comments, which allow anyone with a github account to provide feedback. For example, if the developer needs more information to be able to reproduce a bug, he or she can simply request it in a comment.

Additional elements of issues are (i) color-coded labels that help to categorize and filter issues, (ii) milestones, and (iii) one assignee responsible for working on the issue. They help developers to filter and prioritise tasks and turn issue tracker into a planning tool for their project.

It is also possible for repository administrators to create issue and pull request (see Rule 3) templates (https://help.github.com/articles/helping-people-contribute-to-your-project/) to customize and standardize the information to be included when contributors open issues. GitHub issues are thus dynamic, and they pose a low entry barrier for users to report bugs and request features. A well-organized and tagged issue tracker helps new contributors and users to understand a project more deeply. As an example, one issue in the OpenMS repository (https://github.com/OpenMS/OpenMS/issues/1095) allowed the interaction of eight developers and attracted more than hundred comments. Contributors can add figures, comments, and references to other *issues* and *pull requests* in the repository, as well as direct references to code.

As another illustration of *issues* and their generic and wide application, we (https://github.com/ypriverol/github-paper/issues) and others (https://github.com/ropensci/RNeXML/issues/121) used GitHub issues to discuss and comment changes in manuscripts and address reviewers’ comments.

## Rule 8. Make your code easily citable, and cite source code!

It is a good research practice to ensure permanent and unambiguous identifiers for citable items like articles, datasets, or biological entities such as proteins, genes and metabolites (see also Box 3). Digital Object Identifiers (DOIs) have been used for many years as unique and unambiguous identifiers for enabling the citation of scientific publications. More recently, a trend has started to mint DOIs for other types of scientific products such as datasets [12] and training materials (for example [13]). A key motivation for this is to build a framework for giving scientists broader credit for their work [14, 15], while simultaneously supporting clearer, more persistent ways to cite and track it. Helping to drive this change are funding agencies such as the NIH (National Institutes of Health) and NSF (National Science Foundation) in the USA and Research Councils in the UK, who are increasingly recognizing the importance of research products such as publicly available datasets and software.

A common issue with software is that it normally evolves at a different speed than text published in the scientific literature. In fact, it is common to find software having novel features and functionality that were not described in the original publication. GitHub now integrates with archiving services such as Zenodo (https://zenodo.org/) and Figshare (https://figshare.com/), enabling DOIs to be assigned to code repositories. The procedure is relatively straightforward (see https://guides.github.com/activities/citable-code/), requiring only the provision of metadata and a series of administrative steps. By default, Zenodo creates an archive of a repository each time a new release is created in GitHub, ensuring the cited code remains up to date. Once the DOI has been assigned, it can be added to literature information resources such as Europe PubMed Central [16].

As already mentioned in the introduction, reproducibility of scientific claims should be enabled by providing the software, the datasets and the process leading to interpretable results that were used in a particular study. As much as possible, publications should highlight that the code is freely available in, for example, GitHub, together with any other relevant outputs that may have been deposited. In our experience, this openness substantially increases the chances of getting the paper accepted for publication. Journal editors and reviewers receive the opportunity to reproduce findings during the manuscript review process, increasing confidence in the reported results. In addition, once the paper is published, your work can be reproduced by other members of the scientific community, which can increase citations and foster opportunities for further discussion and collaboration.

The availability of a public repository containing the source code does not make the software open source *per se*. You should use an OSI approved license (https://opensource.org/licenses/alphabetical) that defines how the software can be freely used, modified and shared. Common licenses such as those listed on http://choosealicense.com are preferred. Note that the LICENSE file in the repository should be a plain-text file containing the contents of an OSI approved license, not just a reference to the license.

## Rule 9. Promote and discuss your projects: web page and more

The traditional way to promote scientific software is by publishing an associated paper in the peer-reviewed scientific literature, though as pointed out by Buckheir and Donoho, this is just advertizing [17]. Additional steps can boost the visibility of a organization. For example, GitHub *Pages* are simple websites freely hosted by GitHub. Users can create and host blog websites, help pages, manuals, tutorials and websites related to specific projects. *Pages* comes with a powerful static site generator called Jekyll (https://jekyllrb.com) that can be integrated with other frameworks such as Bootstrap (http://getbootstrap.com/) or platforms such as Disqus (https://disqus.com/), to support and moderate comments.

In addition, several real-time communication platforms have been integrated with GitHub such as Gitter (http://gitter.im) and Slack (https://slack.com/). Real-time communication systems allow the user community, developers and project collaborators to exchange ideas and issues, and to report bugs or get support. For example, Gitter is a GitHub-based chat tool that enables developers and users to share aspects of their work. Gitter inherits the network of social groups operating around GitHub repositories, organizations, and issues. It relies on identities within GitHub, creating IRC (Internet Relay Chat)-like chat rooms for public and private projects. Within a Gitter chat, members can reference issues, comments, and pull requests. GitHub also supports wikis (which are version-controlled repositories themselves) for each repository, where users can create and edit pages for documentation, examples, or general support.

A different service is Gist (https://gist.github.com), which represents a unique way to share *code snippets*, single files, parts of files, or full applications. Gists can be generated in two different ways: public *gists* that can be browsed and searched through *Discover* (https://gist.github.com/discover), and secret gists that are hidden from search engines. One of the main features of Gist is the possibility of embedding code snippets in other applications, enabling users to embed gists in any text field that supports JavaScript.

## Rule 10. Use GitHub to be social: follow and watch

In the same way as researchers are following developments in their field, scientific programmers could follow publicly available projects that might benefit their research. GitHub enables this functionality by *following* other GitHub users (see also Rule 2) or *watching* the activity of projects, which is a common feature in many social media platforms. Take advantage of it as much as possible!

## Conclusions

If you are involved in scientific research and have not used git and GitHub before, we recommend that you to explore its potential as soon as possible. As with many tools a learning curve lays ahead, but several basic yet powerful features are accessible even to the beginner and may be applied to many different use-cases [6]. We anticipate the reward will be worth your effort. To conclude, we would like to recommend some examples of bioinformatics repositories in GitHub (Table 1) and some useful training materials, including workshops, online courses and manuscripts (Table 2).

## Acknowledgments

The authors would like to thank C. Titus Brown for mentioning the manuscript on social media, leading to additional contributions and further improvements. We also thank Peter Cock (peterjc) for helpful suggestion contributed directly though GitHub.

Y.P.R is supported by the BBSRC PROCESS grant (reference BB/K01997X/1) and by the BBSRC Quantitative Proteomics grant (reference BB/I00095X/1). R.W. is also funded by grant BB/I00095X/1. J.A.V. is supported by the Wellcome Trust (grant number WT101477MA). J.U. and T.S. are funded by the BMBF grant de.NBI - German Network for Bioinformatics Infrastructure (FKZ031 A 534A and FKZ031 A 535A). L.G. is supported by the BBSRC Strategic Longer and Larger grant (Award BB/L002817/1). F.V.L. is supported by NIH grant number R01-GM-094231. A.K. is supported by the EPSRC Collaborative Computational Project CoDiMa (reference EP/M022641/1). R.M.F. is supported by NSF grant number 1252893. T.P. is supported by the National Institutes of Health through grant R01-EB-017205. K.B. is funded by the Novo Nordisk Foundation.

## Supporting Information Legends

Supplementary File S1. Supplementary Information including three sections: Git Large File Storage (LFS), Testing Levels of the Source Code and Continuous integration, and Source code documentation.

